# Suboptimal proteome allocation during changing environments constrains bacterial response and growth recovery

**DOI:** 10.1101/2021.04.28.441780

**Authors:** Rohan Balakrishnan, Terence Hwa, Jonas Cremer

**Author notes:** Please address correspondence to JC and RB.

## Abstract

To sustain growth in fluctuating environments microbial organisms must respond appropriately. The response generally requires the synthesis of novel proteins, but this synthesis can be impeded due to the depletion of biosynthetic precursors when growth conditions vary. Microbes must thus devise effective response strategies to manage depleting precursors. To better understand these strategies, we here investigate the active response of *Escherichia coli* to changes in nutrient conditions, connecting transient gene-expression behavior to growth phenotypes. By synthetically modifying the gene expression during changing growth conditions, we show how the competition by genes for the limited protein synthesis capacity constrains the cellular response. Despite this constraint, cells substantially express genes that are not required, severely slowing down the response. These findings highlight that cells do not optimize growth and recovery in every encountered environment but rather exhibit hardwired response strategies that may have evolved to promote growth and fitness in their native environment and include the regulation of multiple genes. The constraint and the suboptimality of the cellular response uncovered in this study provides a conceptual framework relevant for many research applications, from the prediction of evolution and adaptation to the improvement of gene circuits in biotechnology.

## Main text

Changing environmental conditions are a hallmark of microbial habitats and microbes have to respond appropriately to thrive^1–6^. For instance, the depletion of a preferred carbon source requires the efficient transitioning to the consumption of another carbon source^7,8^. Several studies have characterized the response to such diauxic shifts by identifying the up- and down-regulation of hundreds of genes^9–11^, and implicating major regulators such as cAMP^12–15^ and ppGpp^16,17^. Executing these different processes is a major challenge for the cell, especially when biosynthetic precursor levels drop during shifts, yet how cells navigate these challenges and strategize an optimal response remains poorly understood. To decipher the fundamental principles shaping the cellular response and growth-kinetics, we here present a quantitative study on cell-physiology connecting gene-expression to growth-phenotypes. We first studied the shift from growth in glucose to growth in acetate, a shift previously used to study growth-transitions^10,18–20^. We grew *E.coli*-K12 cells in batch cultures and tracked growth by measuring optical density (**Fig. 1A**). The shift from growth on glucose (blue zone) to acetate (red zone) is accompanied by a period of growth arrest (lag-time *τ_lag_*) lasting ~3.5h (gray zone). The lag is also illustrated by the drop of the instantaneous growth-rate during the shift (**Fig. S1A**). From the metabolic perspective, this transition requires a switch from glycolytic pathways to the activation of the glyoxylate shunt and gluconeogenesis pathways^21,10,22,18^ so that the synthesis of amino acids and other growth precursors (green arrows) can continue (**Fig. 1B** and **Fig. S1B**). Hence, following glucose depletion, the synthesis of the glyoxylate shunt enzymes (AceB, AceA) and gluconeogenesis enzymes (MaeA, MaeB, Pck, PpsA) are required before growth can resume on acetate (**Fig. S1B**). But why does it take so long for the few required enzyme types to reach sufficient concentrations for growth to resume?

**Fig. 1:**
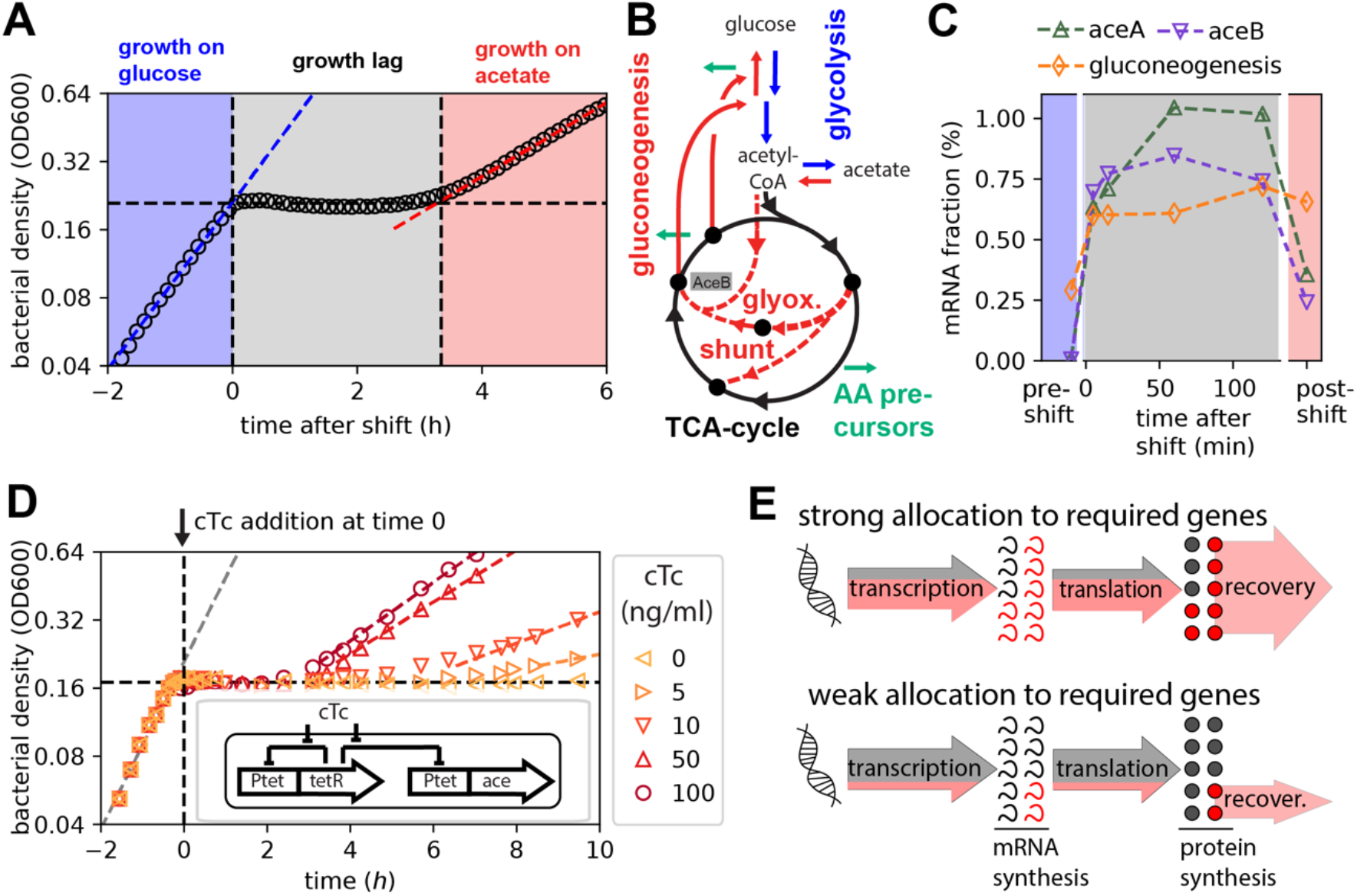
Diauxic shift from glucose to acetate. **(A)** Diauxic growth of WT *E. coli* (NCM3722) in minimal media containing glucose and acetate, bacterial density measured as optical density (OD600). Growth on glucose is captured by the exponential fit (blue dashed line), and proceeds until glucose runs out at time = 0 h. This is followed by a period of growth-lag lasting ~3.5h before exponential growth resumes on acetate (red dashed line). **(B)** The central carbon metabolism pathways are illustrated along with the nodes branching out into amino acid precursor synthesis (green arrows). Glucose to acetate diauxie requires switching from the pathways facilitating glucose consumption (represented in blue) to those responsible for acetate consumption (represented in red). More details in **Fig. S1B**. **(C)** The mRNA fractional abundances for *aceB*, *aceA* and the gluconeogenesis genes (summed abundance of *maeA, maeB, pck,* and *ppsA* genes) were estimated by RNA sequencing performed at various time points during the diauxic transition as indicated on the x-axis. **(D)** Lag-times for controlled titration of *aceBA* expression using an inducer construct in strain NQ1350 (inset). Addition of chlorotetracycline (cTc) removes the tetR repression and induces *aceBA* expression. Increased expression of *aceB/aceA* during the response (inducer added at time = 0h) shortens the lag-time. **(E)** Of the overall transcription and translation fluxes (arrows), a strong allocation towards the expression of shunt and gluconeogenesis genes (red) increases the novel synthesis of required enzymes and should thus lead to faster growth recovery; a weak allocation towards these genes should lead to slower recovery.

To characterize the time-dependence of gene expression, we determined mRNA abundances at various time-points during the growth transition using RNA-sequencing (RNA-Seq, see **SI Text**). The mRNAs of the glyoxylate shunt and gluconeogenesis genes, represented as fraction of total mRNA, increase immediately (< 5 minutes) following glucose depletion and these increased levels are maintained through the duration of the growth lag (**Figs. 1C**). Given such a rapid regulatory response, the speed at which the transcriptional program changes is likely not the reason for long lag-times, but it is rather the expression strength that could be important. To test this idea, we first employed a strain in which the native promoter of the *aceBAK* operon is replaced by the titratable promoter P_*tet*_^20^. Lag times are greatly reduced when increasing amounts of the inducer chlorotetracycline (cTc) are added at the moment of glucose depletion (**Figure 1D**). Since the *aceBA* expression levels prior to glucose depletion are unperturbed, and thus uniform among the cultures, the decrease of lag times with increasing cTc concentrations highlights the significance of the active response to changing conditions in determining the transition kinetics. Following these results we wondered if lag-times emerge due to a fundamental competition for shared resources such as RNA polymerase and ribosomal activity which could be particularly limited during the shift: If a larger portion of the limited transcriptional and translational fluxes are allocated to the synthesis of the required mRNAs and proteins (**Fig 1E top**), the shunt and gluconeogenesis enzymes become available to replenish precursors earlier than in the case with a lower allocation of resources towards these genes (**Fig 1E bottom**).

To probe this allocation picture, we next employed a titration construct to over-express *lacZ*^23^, the product of which hydrolyses lactose and is thus useless for growth in glucose and acetate (**Fig. 2A**, inset): when transcriptional and translational resources are diverted towards LacZ synthesis during the response to changing conditions, the protein itself adds no benefit to the cell and thus acts as a sink for shared resources which should extend lag-times. In line with this expectation, when inducing *lacZ* expression by adding various levels of the inducer chlortetracycline (cTc) at the moment of glucose depletion, we observed that lag-times increase strongly from *τ_lag_* = 3.9 *h* at 0 ng/ml cTc to *τ_lag_* = 12.2 *h* at 7 ng/ml cTc (**Figs. 2A & S2A**). To further explore this effect, we measured the mRNA levels of *lacZ* and the required shunt genes *aceB* and *aceA* by qPCR, 10 minutes after the shift. The abundance of *lacZ* mRNA increases with inducer concentration (**Fig. S2B**) in direct relation to the lag-time (**Fig. 2B**). Notably, as *lacZ* mRNA is dialed up, *aceB* and *aceA* expression is reduced (**Fig. 2C, Fig S2CD**), explaining the longer lag-times based on a lower expression of these required enzymes (**Figs. 2D)**. Hence, upon synthetically introducing a resource scarcity during an environmental shift, these observations indeed suggest that the allocation of limited shared resources determines the cellular response and thus lag-times.

**Fig. 2:**
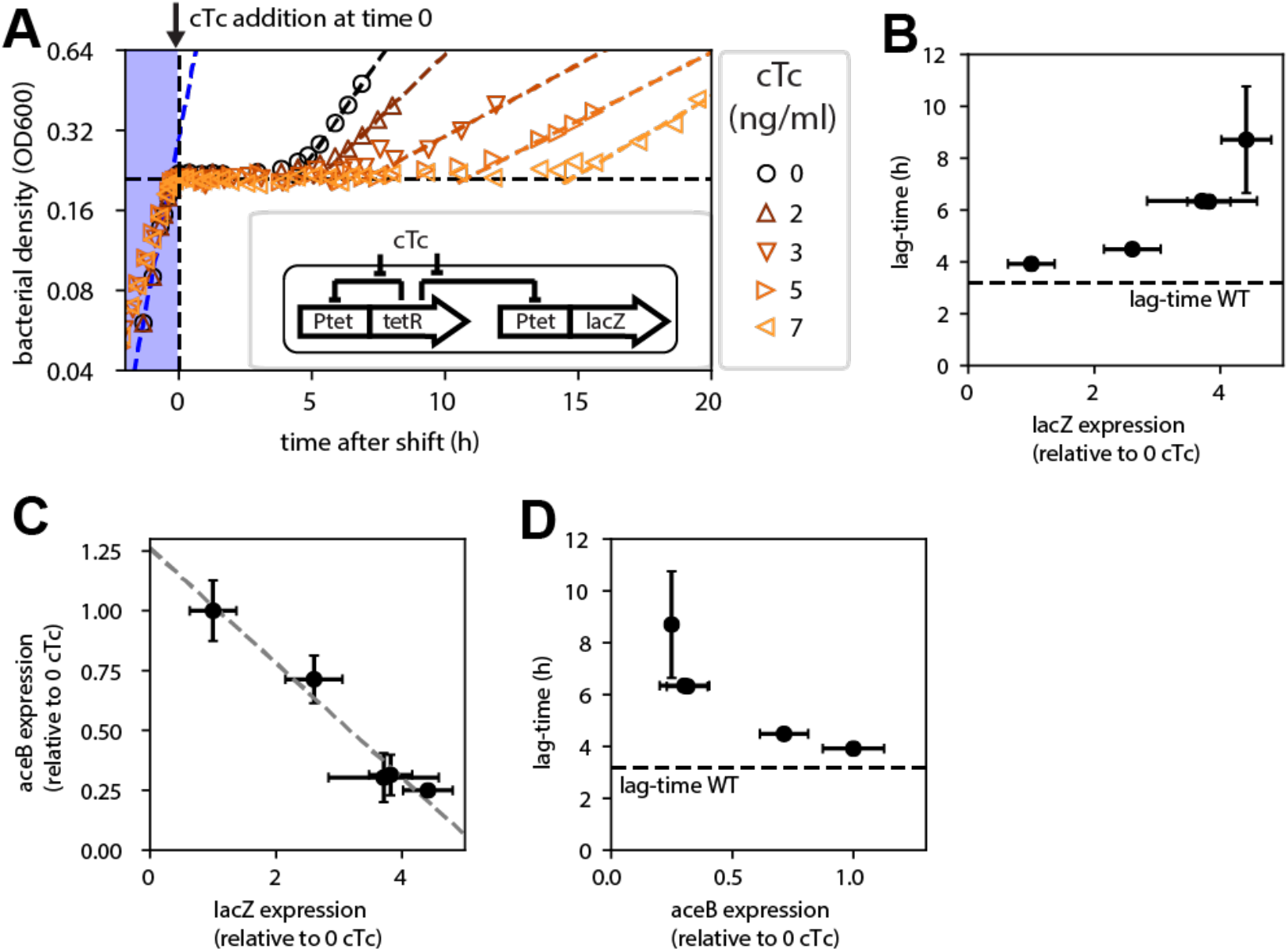
Expression of a non-needed gene inhibits expression of required genes and elongates lag-times. **(A)** A plasmid system (inset) is used to control the expression of the non-required gene *lacZ* using the strain NQ1389. *lacZ* expression was induced using cTc to varying degrees at the moment of glucose depletion (time = 0 h) using the indicated range of cTc concentrations (strain NQ1389). Diauxic growth conditions with glucose and acetate same as in **Fig. 1**. **(B-D)***lacZ* mRNA resulting from the different degrees of induction as well as *aceB* mRNA in the same cultures were measured by qPCR and scaled relative to their corresponding abundances observed in absence of induction. The lag-times observed in panel A are plotted against the respective *lacZ* abundances (B), where the dashed line represents the 3.5h lag observed for the WT strain (no induction, **Fig. 1A**). *lacZ* and *aceB* mRNA levels are inversely related (C), line shows linear fit. *aceB* mRNA abundance is inversely related to the lag-time (D). qPCR and lag-time data plotted as the average of at least three independent replicates with error bars indicating standard errors.

To better understand how the competition for shared transcription and translation resources can have such drastic impacts on growth transitions, we next formulated a kinetic model of growth which focuses on protein synthesis as the most resource demanding process of biomass synthesis. The model builds on recent advances to describe growth^24–29^ and explicitly considers amino acid precursors, their synthesis by metabolic enzymes, and their utilization by ribosomes in form of charged tRNA (**Fig. S3**). A key feature of the model is that only a fraction of the ribosomes synthesize the enzymes (eg. *AceB*) that supply the precursors, while the rest of the translation flux is diverted to the synthesis of other proteins (**Fig 3A**). The consumption of amino acids precursors however depends on the (total) protein synthesis, leading to a feedback between protein synthesis and precursor supply. During the diauxic transition, where there is a sudden depletion of cellular amino acid pools following the runout of the preferred carbon source, this can lead to cells being “trapped” in a low precursor state. The mathematical analysis shows that such states can persist for hours when (i) the required proteins such as AceB have not been synthesized in sufficient numbers yet, and (ii) the remaining amino acid levels are insufficient to support the synthesis of new proteins (**Fig. S4A-E**). A direct way to mitigate this trap is to allocate a larger fraction of the translation flux towards the synthesis of the required enzymes (**Fig. S4F-J**). Accordingly, lag-times fall drastically with a higher allocation towards the synthesis of required enzymes (**Fig. 3B)**, reflecting the lag-time changes observed when overexpressing the required or non-required genes *aceBA* and *lacZ* (**Figs. 1D & 2**). Taken together, our experiments and theoretical analyses establish mechanistically how the allocation of limited resources during the shift shapes growth transitions, outlining a range of possible allocational behaviors with varying consequences on the growth transition kinetics.

**Fig. 3:**
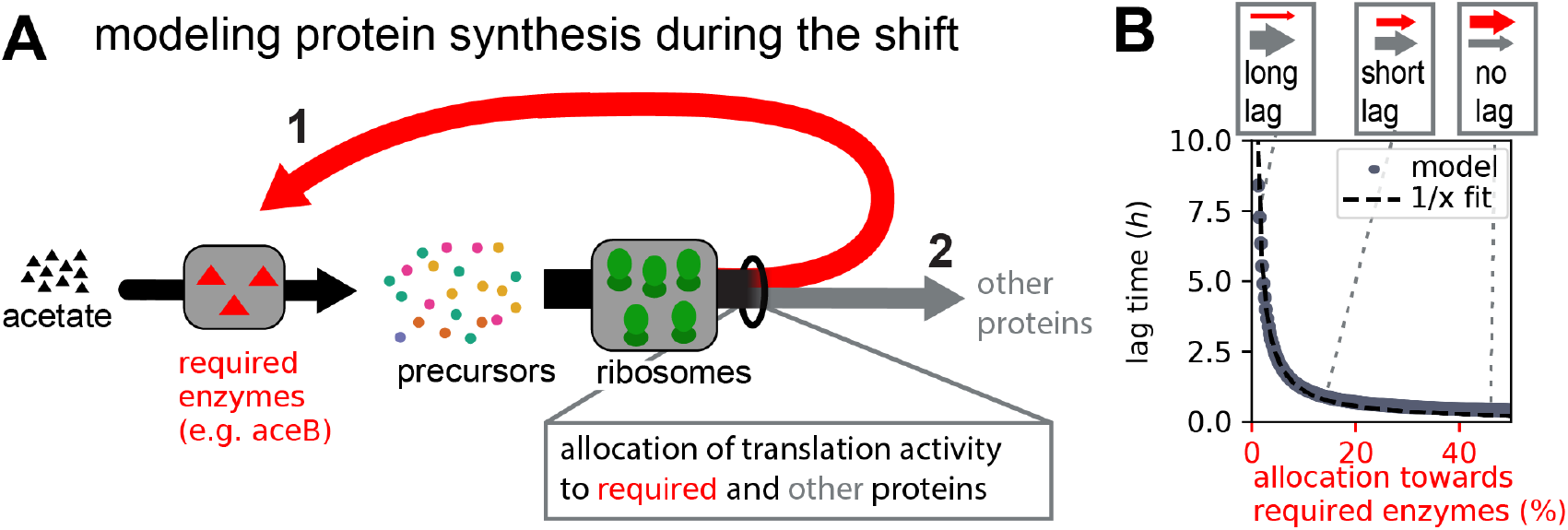
Modeling growth kinetics during the shift. **(A)** Essential dynamics during the shift from growth on glucose to growth on acetate: Protein synthesis by ribosomes depends on the availability of precursors which itself depends on the availability of metabolic enzymes which utilize acetate to provide novel precursors. When ribosomes synthesize more of these required enzymes (red arrow 1) instead of other proteins (gray arrow 2) novel precursors are generated from acetate (black arrow) faster and growth thus resumes faster (**Figs. S3 and S4**). **(B)** Lag-times fall reciprocally with the allocation towards required proteins (the fraction of mRNA encoding for required proteins). Allocation of translation activity towards different proteins are represented by the weight of red and gray arrows (top). Full model introduced in **Fig. S3 and SI Text 2**. Model parameters provided in **Table S3**.

We next ask where in this range of allocational behaviors does native *E. coli* (no synthetic overexpression) fall, and whether the long lag-times observed for WT *E. coli* (**Fig. 1A**) also emerges from the synthesis of non required proteins during the shift. It has long been known that *E. coli* growing steadily on poor carbon sources (such as acetate and glycerol) express several catabolic enzymes, despite the absence of their specific substrates^30,31^. Allocating resources towards such “idling” proteins during growth transitions would map *E. coli* towards the left of the plot in **Fig 3B**, with lag-times substantially larger than those expected with specialized regulatory strategies in which only the required genes are expressed. To quantify the transcriptome fraction that potentially encodes idling proteins, we first analyzed transcriptomics measurements for *E. coli* grown under steady-state conditions with either glucose or acetate as carbon sources. We considered the more abundant half of the genes and determined those genes which are expressed at least 2-fold more in acetate compared to glucose. Most of these genes belong to a few functional categories including transport, motility, and catabolism (**Figs. 4A & S5A**). Furthermore, as seen from our transcriptomics data collected over the course of the growth transition, these genes are upregulated immediately following glucose depletion (**Fig. 4B**). Yet, most of these functions are not expected to be useful for the growth transition to acetate: motility is not needed in shaking environments and most of the uptake transporters are involved in transporting other nutrients besides acetate. The expression of these non-required genes could impede the allocation of shared resources towards the genes encoding for shunt and gluconeogenesis enzymes. To test this idea we next considered the growth behavior of deletion strains lacking motility genes, the category which showed most increased expression in acetate (**Fig. 4A**). We considered two mutants, *ΔfliC* and *ΔflhD*, which either do not express the flagella protein FliC or the motility master regulator FlhDC required for the expression of flagella, motors, and the chemotaxis apparatus^32^. The mRNAs of these genes comprise up to ~15% of the total transcription in the WT strain growing on acetate and could considerably reduce the allocation of resources towards the shunt and gluconeogenesis genes. When grown under steady state conditions with either glucose or acetate as sole carbon sources, we observe that none of these deletion strains have any defect in steady growth rates, but rates even increased substantially for growth on acetate (**Fig. 4C**), supporting the idea that these gene products are indeed useless for growth in the probed conditions. The increase of growth was also observed for other carbon sources besides acetate (**Fig. S6**) and provides direct support for the idea that gene regulation is not optimized for steady state growth^33–35^.

**Fig. 4:**
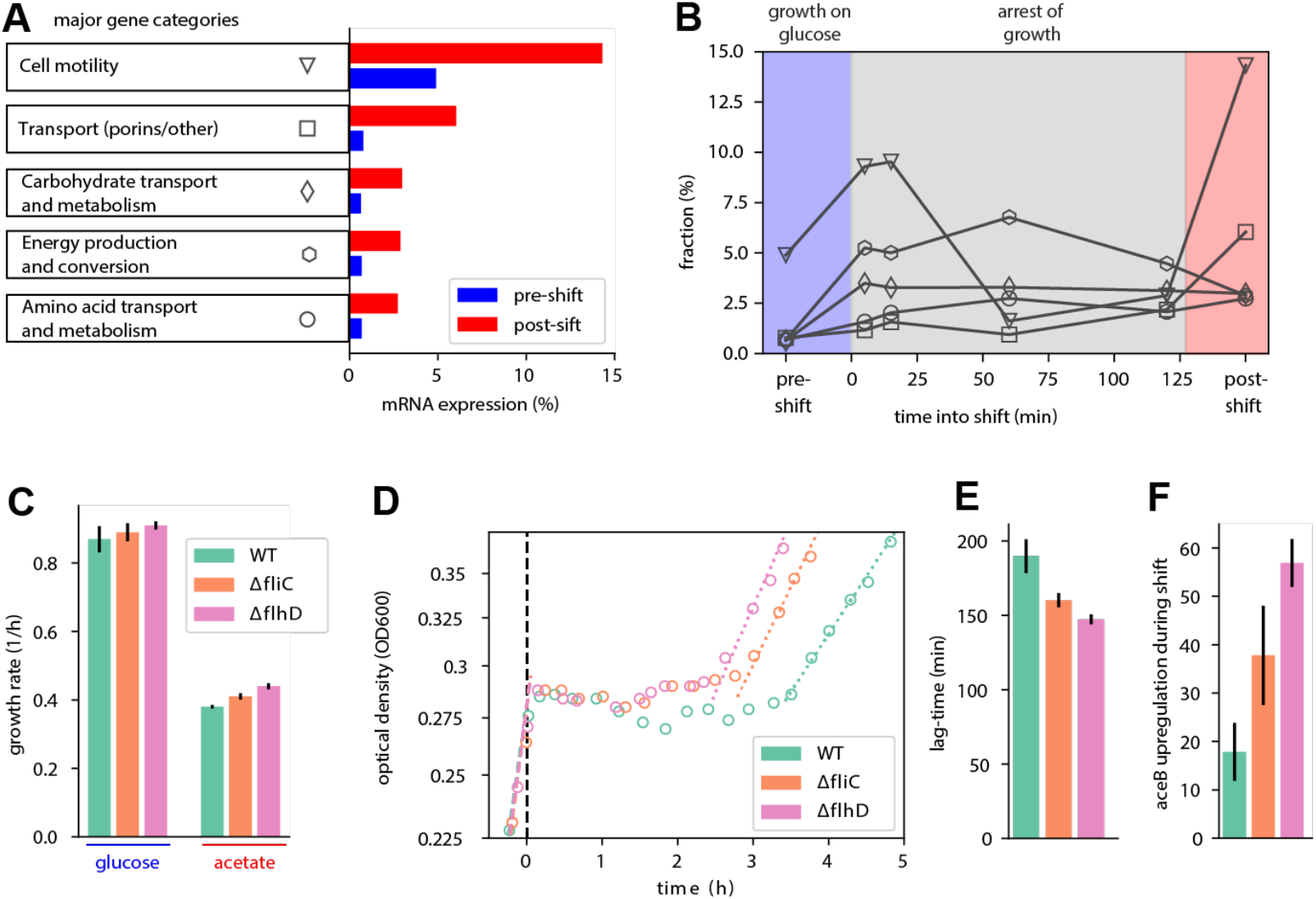
Effect of non-required gene expression on the growth transition. **(A)** The major gene categories up-regulated in balanced growth on acetate (red) compared to that on glucose (blue). mRNA abundances determined by RNA-Seq are represented as percent of total mRNA in the given growth condition. Genes are categorized using the COG classification^43^ along with manual curation of the porins using ecocyc.org. **(B)** For the various gene categories in panel A, the temporal kinetics of expression during the diauxic shift is plotted. Time 0 indicates time when glucose runs out. **(C)** Steady-state growth in glucose and acetate for the WT and the motility deletion strains *ΔfliC* and *Δflh*. Error bars represent standard errors of three replicate experiments. Additional growth conditions are shown in **Fig. S6**. WT (NCM3722) used in (A,B). **(D)** Growth transitions for the motility deletion strains *ΔfliC* and *ΔflhD* (blue and magenta) are substantially faster than that for the WT strain (red). **(E)** Resulting lag-times of the transitions shown in (A). **(F**) Upregulation of the *aceB* gene for the WT and the motility deletion strains during the transition relative to their pre-shift levels. Error bars represent the standard error of two independent replicates.

Given the vast expression of motility genes immediately following glucose depletion (**Fig 4B**), we next probed how the synthesis of these non-required proteins affects the growth transition from glucose to acetate by quantifying lag-times for the two deletion strains. Lag-times are shorter for the deletion strains than for the WT strain **(Fig. 4DE)**, suggesting that the deletion strains are better at directing RNA-polymerases and ribosomes towards the synthesis of the shunt and gluconeogenesis enzymes. To test whether the deletions lead to increased transcription of the required genes, we used qPCR to track the upregulation of the glyoxylate gene *aceB* during the transition. The *ΔfliC* and *ΔflhD* strains indeed show an up to 3-fold higher upregulation of *aceB* compared to the WT (**Fig. 4F**), explaining the reduced lag times based on the higher resource allocation towards the required genes. These results are consistent with the previous observation that the overexpression of non-needed genes leads to lower *aceB* expression levels and thus longer lag times; **Fig. 1**).

Finally, given the results for the diauxic growth on glucose and acetate, we asked whether the expression of idle proteins and their competition with required metabolic enzymes is also responsible for the lag-times observed in other diauxic growth conditions. We tested other conditions originally reported by Monod^7,8^, inducing shifts from glucose to glycerol, xylose, and maltose. These carbon sources enter the central metabolic pathway at different steps (**Fig 5A**) and do not require the flux-reversal from glycolysis to gluconeogenesis which was recently suggested to explain long lags in growth-transitions^14^. Instead, these carbon sources necessitate the synthesis of other unique sets of transporters and catabolic enzymes for growth to resume and are thus distinct from acetate. Using the motility deletion strain, we consistently find reduced lag-times for all transitions (**Fig. 5B-F**). Hence, the allocational constraint of shared protein synthesis resources is a general principle governing a range of different diauxic transitions.

**Fig. 5:**
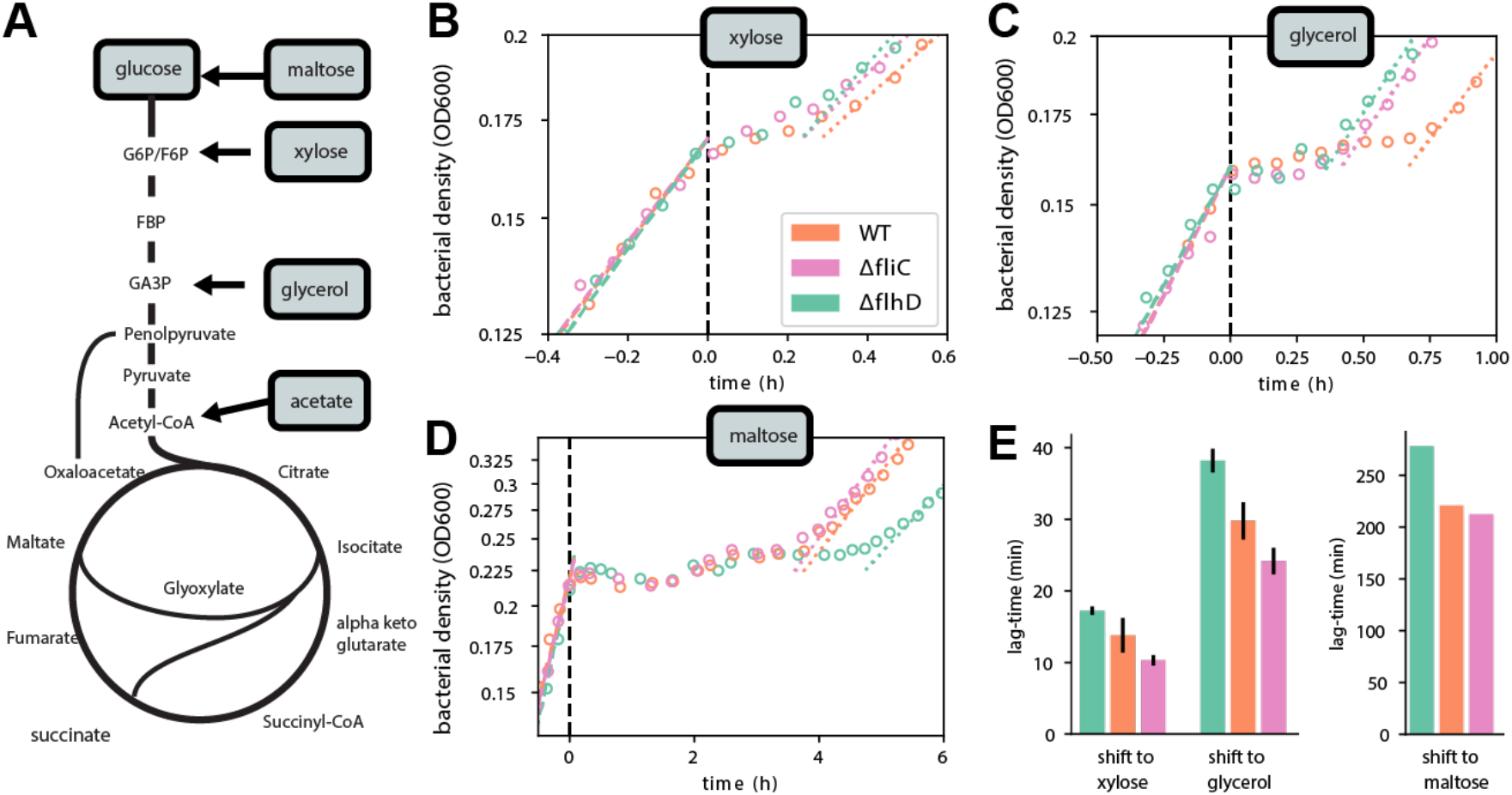
Non-required proteins are responsible for lags during many diauxic shifts. **(A)** The expression of different uptake and metabolic proteins are required for the utilization of different carbon sources. Points of entry into the central metabolism are indicated. **(B-D)** Diauxic growth kinetics for growth on glucose and a different secondary carbon sources for the WT (red) and the motility deletion strains *ΔfliC* and *ΔflhD* strain (blue and magenta). **(E)** Lag-times for the different transitions.

## Discussion

Since the pioneering growth physiology studies by Monod, lag-times have been perceived as the preparation time cells require to adjust to a new environment before growth can resume^1,4,7,8,17,36^. In line with this idea, several studies in bacteria and yeast suggest that the expression of required genes before the nutrient shift can reduce lag-times but can come with the cost of slower growth rates, implicating a tradeoff between lag-times and growth in the pre-shift condition^3,20,37–39^. In this quantitative study, we instead focus on the cellular response to environmental changes. We establish how the competition for shared resources towards novel protein-synthesis fundamentally constrains the cellular response and shape growth-transitions. The duration of growth recovery depends on whether only the required genes (specific response) or a diverse array of genes (diversifying response) are expressed during the response to the environmental change. Tweaking the specificity of the response can substantially vary lag-times.

Our observations thus call for a revision of the previously proposed explanations for growth transitions: While pre-shift growth/lag trade-offs may affect lag-times, growth-transitions are determined first and foremost by a compromise between specific versus diversifying responses during the shift. Mechanistically, this compromise stems from a constraint on the allocation of shared resources towards the synthesis of novel proteins during the response. This constraint opens a multitude of possible response strategies, ranging between extremely specific and diversifying mechanisms. A highly specific response tailored towards the growth conditions encountered can ensure fast growth-transitions across many shifting conditions. But harboring specific regulatory pathways for multiple different nutrient sources can be cumbersome. In addition, a diversifying response involving the expression of a broad array of genes might provide benefits in certain ecological scenarios. One plausible means to enforce the strategic compromise between specific and diversifying responses is a heterogeneous response within the population with strong differences of gene expression and growth across cells. The constraint revealed in this study thus provide a potential strong rationalization for these \ “bet-hedging” stretegies^19,40–42^. However, for the diauxic condition probed in this study, no heterogeneity was observed^20^, suggesting that responses rather arise from hardwired gene regulatory circuits in homogenous populations.

The response of *E. coli* includes the expression of several genes such as diverse transporters and flagella components which may not be required in the encountered environment. The response is thus diversifying instead of specific which may be rationalized from an ecological perspective: Besides glucose, *E. coli* encounters many other sugars and amino acids within the mammalian intestine, and swimming is expected to play a crucial role in the strong flow environment prevalent within the intestine^43,44^. The diversifying response thus appears to be tailored towards coping with the fluctuating environments typical for the gut. But it is exactly such a response that constrains resource allocation towards the specific required genes during other transitions, as those encountered in laboratory experiments, leading to long lag times. Accordingly, strains evolving in typical laboratory environments would be expected to lose their diversifying response. In fact, we observe evidence for this hypothesis embedded in previously published results of the long-term evolution experiment by Lenski and colleagues^45,46^: the evolved strains exhibit loss of growth on many carbon sources and motility^47^. In the light of our findings, it would be interesting to see if the selective advantage for these evolved strains is indeed, in part, due to shorter lag-times. Finally, by describing the mechanistic constraints shaping the cellular response, our work establishes the physiological framework for rational strain engineering in biotechnological applications: By trimming down the diverse response highly optimized behaviors such as short lags and high yields can be instilled.

## Acknowledgement

We thank members of the Cremer and Hwa research groups for discussions. Part of the transcriptomics sequencing work was performed at the Institute of Genomic Medicine, University of California, San Diego.

## Supplementary Figures

**Fig. S1:**
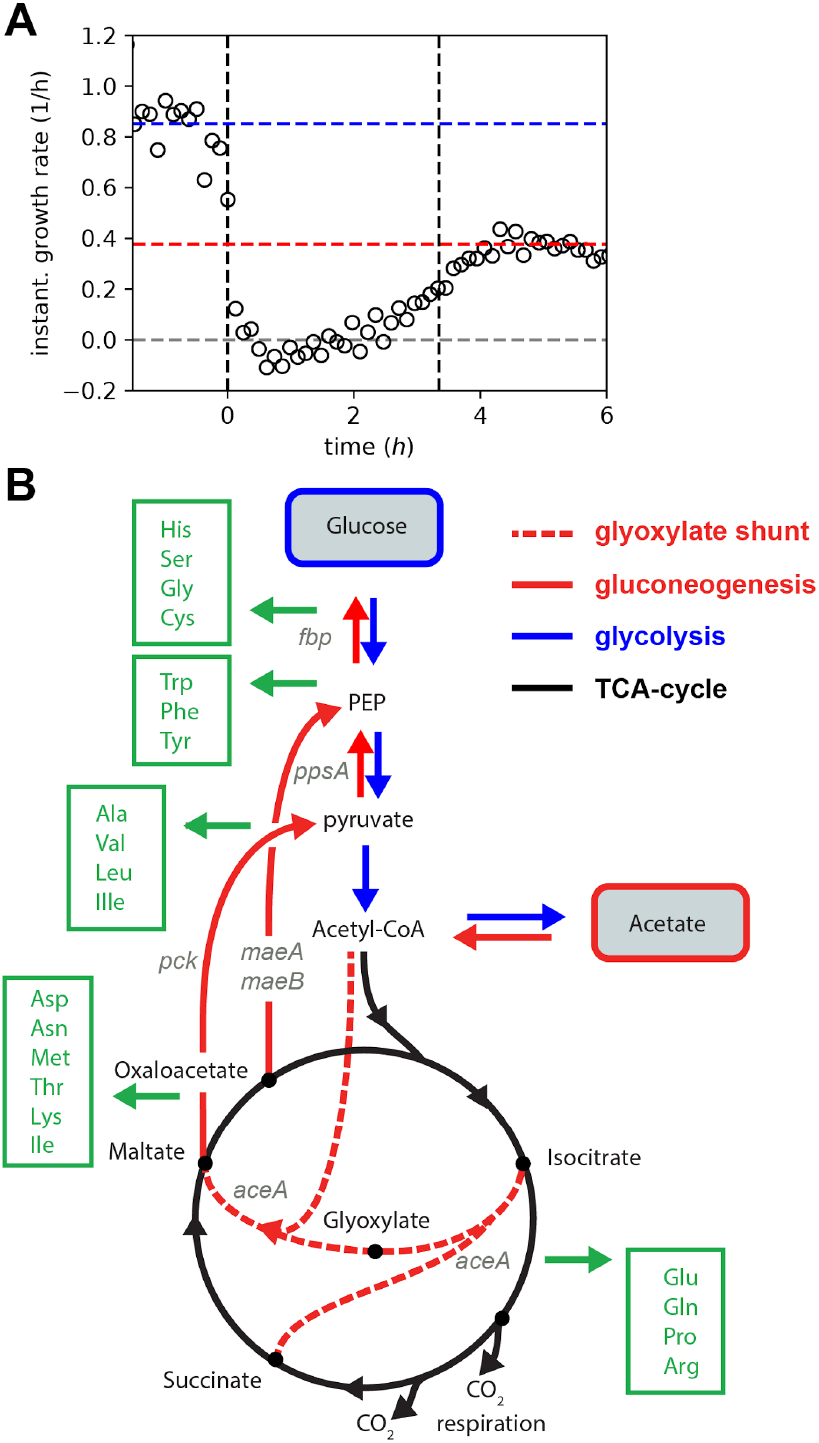
Diauxic growth on glucose and acetate: transition kinetics and metabolic requirements. **(A)** Instantaneous growth-rate of the WT (NCM3722) is derived by calculating the derivative of the growth-rate divided by the optical density (growth curve shown in **Fig. 1A**). Cells first consume glucose and grow exponentially at a rate of ~0.9/h (blue horizontal line). Following glucose depletion (defined as time 0), the instantaneous growth rate immediately falls and growth stops. After a phase of no growth, growth gradually begins to approach a rate of 0.4/h, the steady state growth rate for growth on acetate (red horizontal line). **(B)** To resume growth on acetate following glucose depletion the supply of amino acids, the major precursors required for biomass synthesis, must be re-established. The various nodes along the central carbon metabolism that branch into the synthesis of different amino acids are indicated in green. For the successful recycling of the 2 carbon molecules per acetate into amino acid precursors, the most essential step is the activation of the glyoxylate shunt^22^: To prevent the loss of the 2 CO2 molecules occurring along the TCA cycle, the carbon flux has to be bypassed. Instead of being converted to alpha-KG, isocitrate is split into succinate and glyoxylate by the enzyme isocitrate lyase (*AceA*). Glyoxylate is then converted to malate by the malate synthase (*AceB*). Succinate and malate generated as a result of the shunt subsequently fuel gluconeogenesis (*MaeA*, *MaeB*, *Pck*, *PpsA*), making available the carbon precursors required for the synthesis of many amino acids (green arrows). Besides amino acids as precursors, protein synthesis also requires substantial amounts of energy, primarily to charge tRNA and drive translation. However, energy supply is unlikely to be the major bottleneck during this shift since cells already express and utilize TCA enzymes during the pre-shift growth which can thus ensure a continuous production of ATP over the course of the transition.

**Fig. S2:**
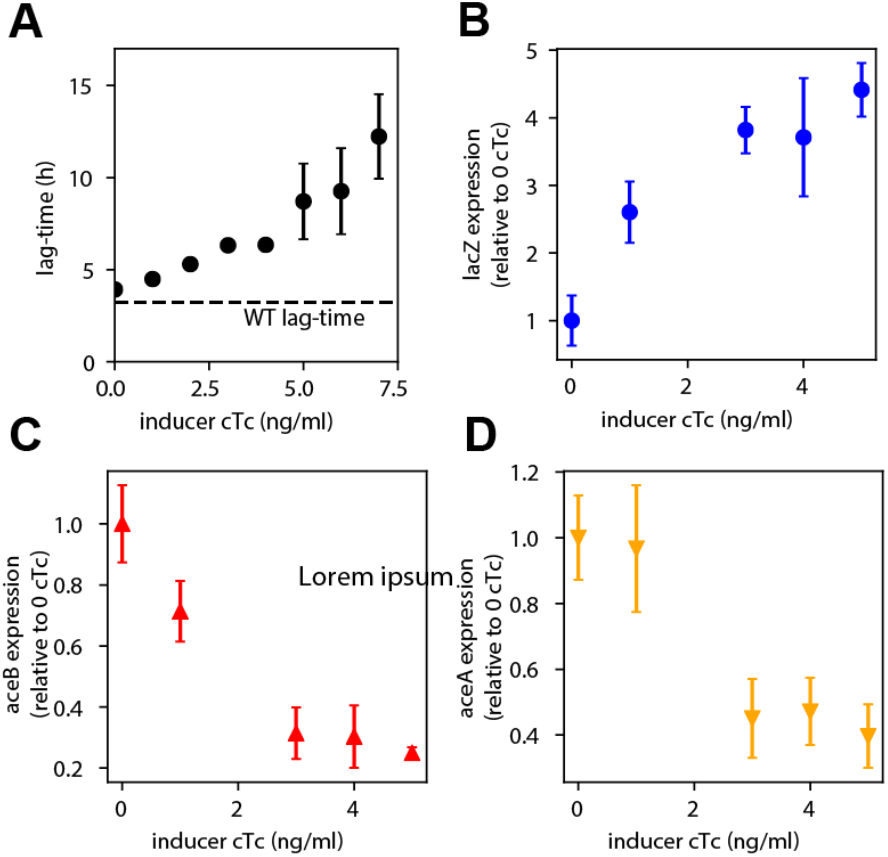
Lag-times and gene expression when overexpressing lacZ. **(A)** Increase in lag-times with increasing inducer levels (strain **NQ1389**). The inducer chlortetracycline (cTc) is added when glucose runs out (**Fig. 2A,** time=0). Dashed horizontal line indicates lag-time for the WT strain (NCM3722). **(B,C,D)** mRNA levels of *lacZ*, *aceB*, and *aceA* at different cTc levels is quantified by qPCR 10 min after the depletion of glucose. mRNA levels of each gene are normalized to the 16S rRNA level, which is known to remain constant during the lag phase^26,48^. These normalized expression levels are shown relative to the expression level in the absence of induction (0 cTc). Error bars indicate standard error for N=3 biological repeats.

**Fig. S3:**
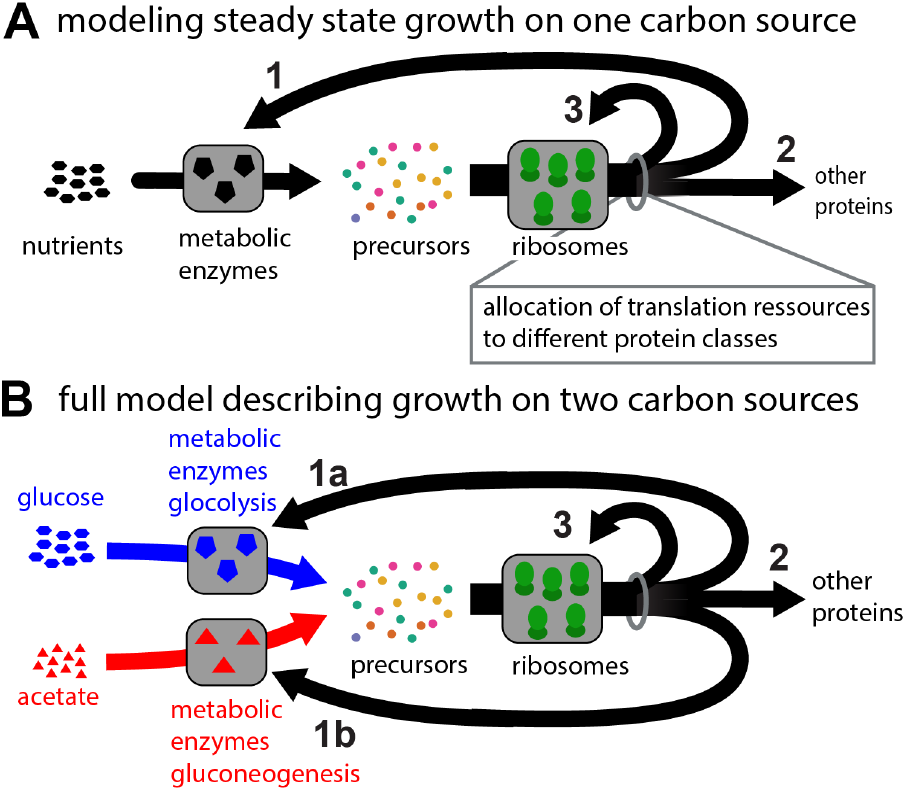
Modeling growth in changing environments. To model growth during transitions, we here build on replicator/allocation models which have been established previously for growth in steady state conditions and for specific shifts. Replicator/allocation models consider protein synthesis by ribosomes towards different proteins^15^ and central to their approach is the allocation of ribosome activity towards the synthesis of different proteins. To illustrate the concept, we here consider steady growth on one carbon source first (**A**). The pool of ribosomes is allocated towards different protein classes such as the proteins for novel ribosomes (A, arrow 3), metabolic proteins needed to provide the precursors when utilizing the carbon source (A, arrow 1) and other proteins (A, arrow 2). Growth depends on the availability of nutrients and how the ribosomal activity is allocated to the different protein classes: high growth rates are achieved with allocation ratios that balance precursor influx provided by the metabolic enzymes and their utilization by ribosomes, such that as many ribosomes as possible can translate at maximum speed. This logic is formulated mathematically in **SI Text 2.1**. Notably, for *E. coli* the allocation of ribosomal activity towards the synthesis of new ribosomes follows indeed a close to optimal regulation scheme preventing the synthesis of idle ribosomes (as manifested in the ribosome content changing with growth rates). To describe growth during shifts we extend this modeling approach and explicitly consider glucose and acetate as two nutrient sources **(B).** In this case, two metabolic protein classes (arrows 1a & 1b) which provide the precursors when cells grow on the two carbon sources (blue and red arrows) and perform glycolysis (blue) or gluconeogenesis (red) respectively. As such, this model structure shares similarities with recent modeling approaches to describe growth during nutrient shifts^24,26,27^. However, our approach is distinguished from previous studies by the inclusion of two key aspects, which are central to the cellular response during growth shifts: i. We consider the highly responsive regulation of transcription and integrate our transcription measurements (**Fig. S5**) which quantifies the immediate expression response of the cell during the shift. ii. we explicitly vary the allocation towards other proteins shift (arrow 2) during the shift, thereby bringing the focus of the study to the allocational constraints acting during the response itself. Notably, right after the shift the model simplifies to the scenario shown in **Fig. 3A** and the consideration of metabolic enzymes (arrows 1 & 2) together with the precursors required to drive novel protein synthesis is sufficient to investigate how long lag times can emerge and how they relate to the expression of non-required proteins. The mathematical formulation of the full model and additional context is provided in **SI Text 2** and results for the switch from growth on glucose to growth on acetate are provided in **Figs. S4**. The central model output, the curve describing how lag-times decrease with an increasing allocation to required proteins, is shown in **Fig. 3B**.

**S4.**
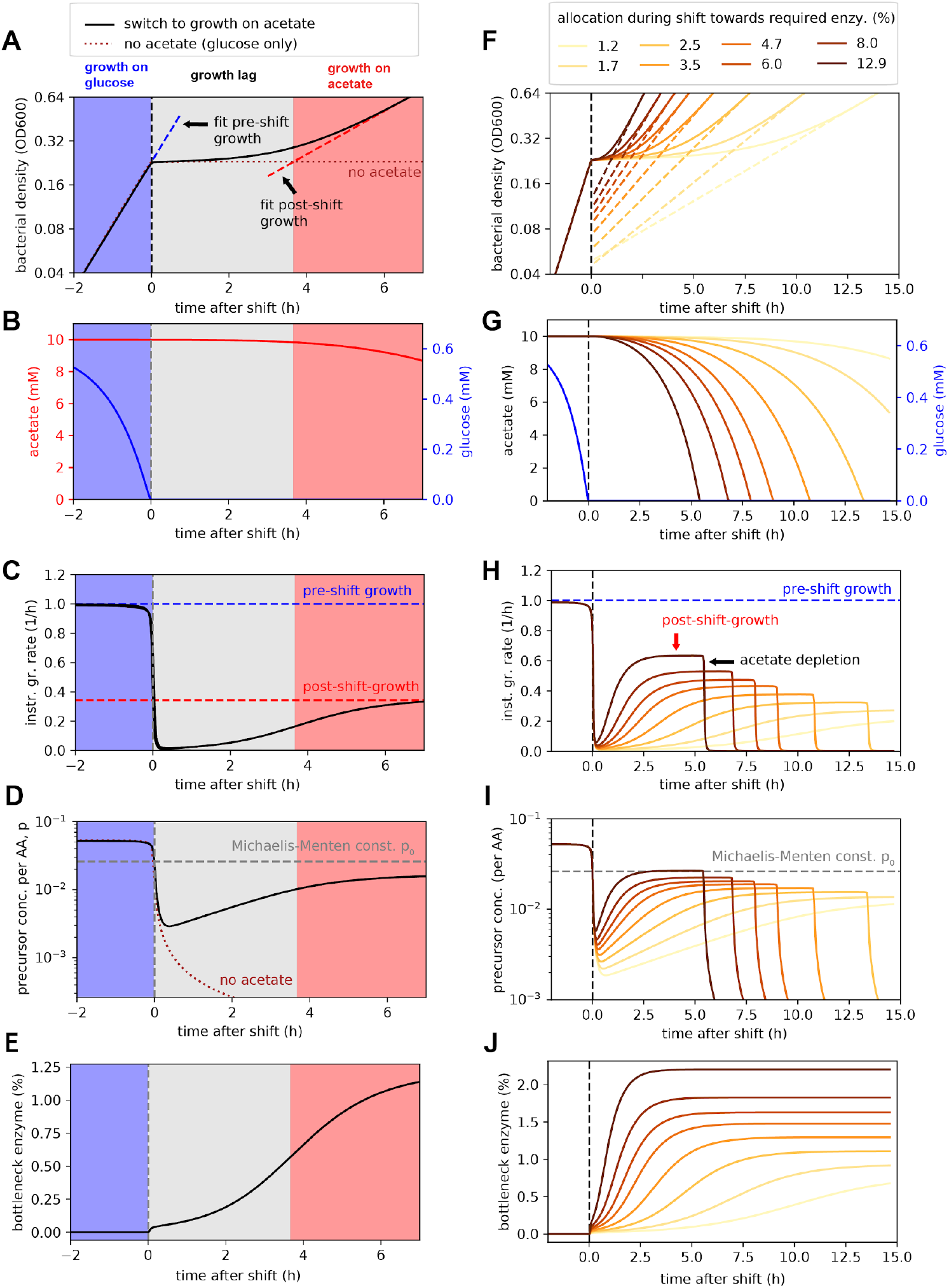
Modeling growth transitions and the consequence of non-required protein expression. Based on our model of growth transitions (**Fig. S3, SI Text 2**) we analyzed growth kinetics during the transition from growth on glucose to growth on acetate. **(A-E)** Change of major model variables (cell density, nutrients, precursors, and required metabolic enzymes) during the shift for a reference condition with a specific allocation towards the metabolic enzymes required for growth on acetate. Growth and nutrient abundance: When glucose is available, cells grow fast (A, blue region) consuming only glucose, and not acetate (B, blue region). Following glucose depletion, precursors fall dramatically (D, gray region) and growth thus stops (A & C, gray region). Cells then only slowly synthesize the required metabolic enzymes like AceB (E) which then slowly support higher precursor concentrations (D, gray region). The case where no metabolic proteins for growth on acetate are synthesized is shown for comparison (D, dashed line); precursor concentrations continue to fall. Growth only recovers to post-shift growth-rates (A & C, red regions) once precursors reach concentrations comparable again to the Michaelis Menten constant (D, dashed horizontal line) which describes the concentration of precursors above which ribosomes can work efficiently with close to maximum translation rates. The growth recovery is slow and spans several hours since cells are trapped in a state where precursor concentrations and the abundance of the metabolic enzymes which can generate new precursors are both low. Accordingly, novel metabolic proteins can only be generated slowly and precursor concentrations thus also recover slowly. **(F-J)** Effect on changing allocation towards the metabolic enzymes required for growth on acetate. Plots show the same variables as in (A-E) but for different allocational behaviour towards the synthesis of novel proteins required during the shift (model parameter*α_Mb,ace,max_*describing the allocation of overall transcription (mRNA fraction) to the required enzymes, see color legend). A higher allocation towards the metabolic enzymes (darker colors) substantially decreases the lag during the growth transition as it leads to much faster accumulation of required metabolic enzymes (J), preventing a dramatic decrease of precursor concentrations right after the shift (I). This thus also leads to a much faster recovery of growth (I). Growth eventually stops once acetate is also consumed (G,H). The change in lag-times for different allocations towards the required metabolic enzymes is shown in **Fig. 3B**. All times shown are relative to the time-point where glucose is depleted and colored regions in (A-F) indicate different growth phases as defined by the intersection of exponential growth curves (dashed lines in A & F) and the density values during the shift, following what was done for the experiments (**Figs. 1A & S1A**). Model parameters as listed in **Table S3**.

**Figure S5:**
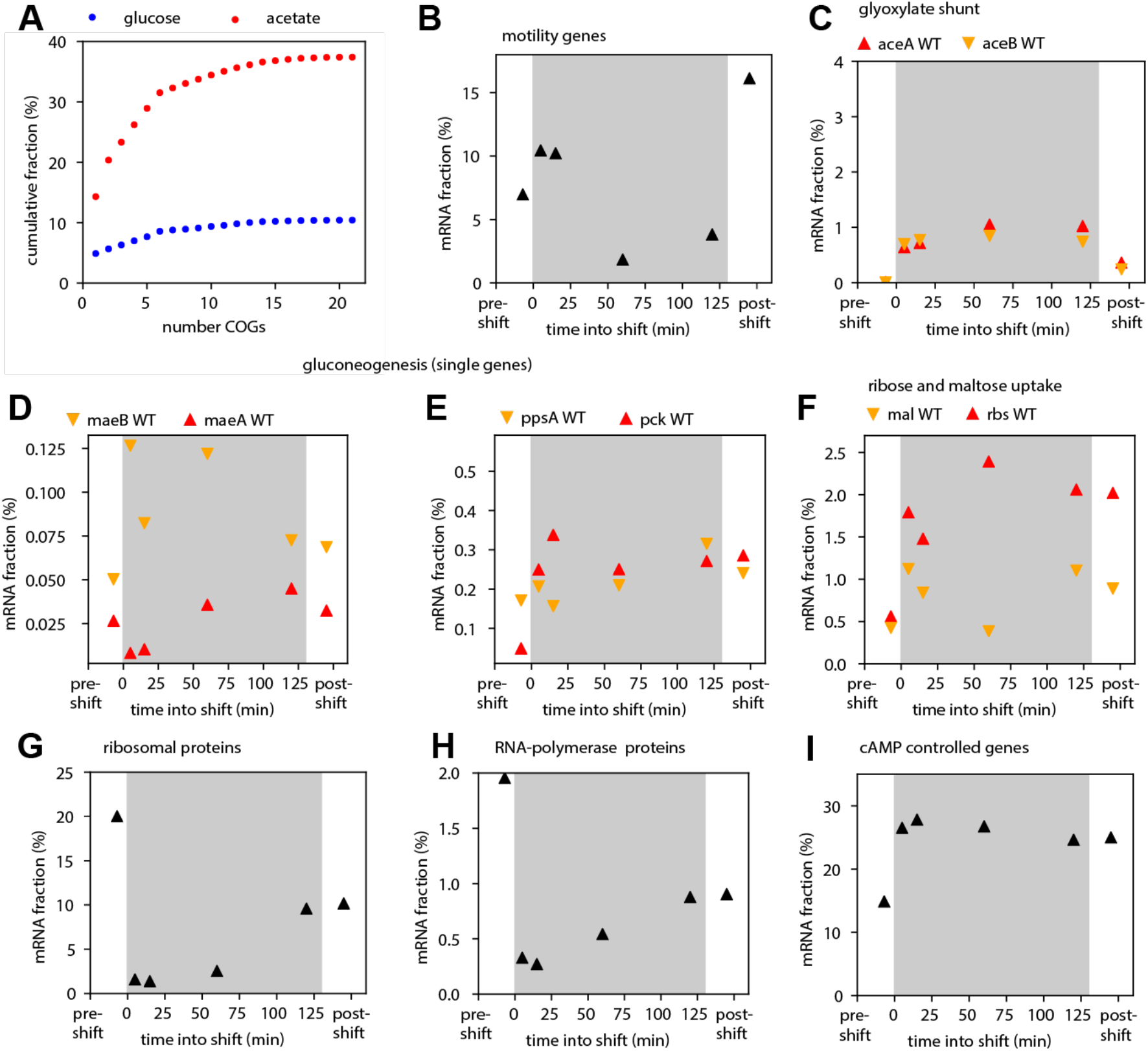
Gene expression during the transition. **(A)** Fractions of genes upregulated in acetate (red) compared to glucose (blue), shown as cumulative sum over most expressed functional groups. Data is derived from transcriptomics measurements using adjusted clusters of orthologous groups (COGs). The 5 most expressed functional groups, shown in **Fig. 4B** during the shift, cover most of the genes which are upregulated. **(B-I)** mRNA abundance for different genes and gene groups during the diauxic shift from glucose (pre-shift) to acetate (post-shift), using RNA-Seq measurements. Measurements for the WT strain NCM3722. **(B)** Motility genes (defined in (10), consisting of *fliC* encoding the major structural component of the flagella and different motor genes) are heavily expressed during the shift. **(C)***rbs* and *mal* genes (detailed version of **Fig. 5C**) are upregulated during the shift. **(D)** Glyoxylate shunt genes (detailed version of **Fig. 5D**) are upregulated. **(E, F)** The upregulation of different gluconeogenesis genes. **(G,H)** Expression of ribosomal protein (summed over all *rpl*, *rpf*, and *rpm* genes) and RNA polymerase genes (*rpoA,B,C*) during the shift. Within 5 minutes these genes are repressed, confirming a responsive down-regulation mechanism. **(I)** The expression of genes known to be under the control of the second messenger cAMP_45_. This group of genes is upregulated during the shift. All data is derived from the same RNA-Seq dataset taking the relative fraction of different genes (and their sum when groups of genes are considered).

**Figure S6:**
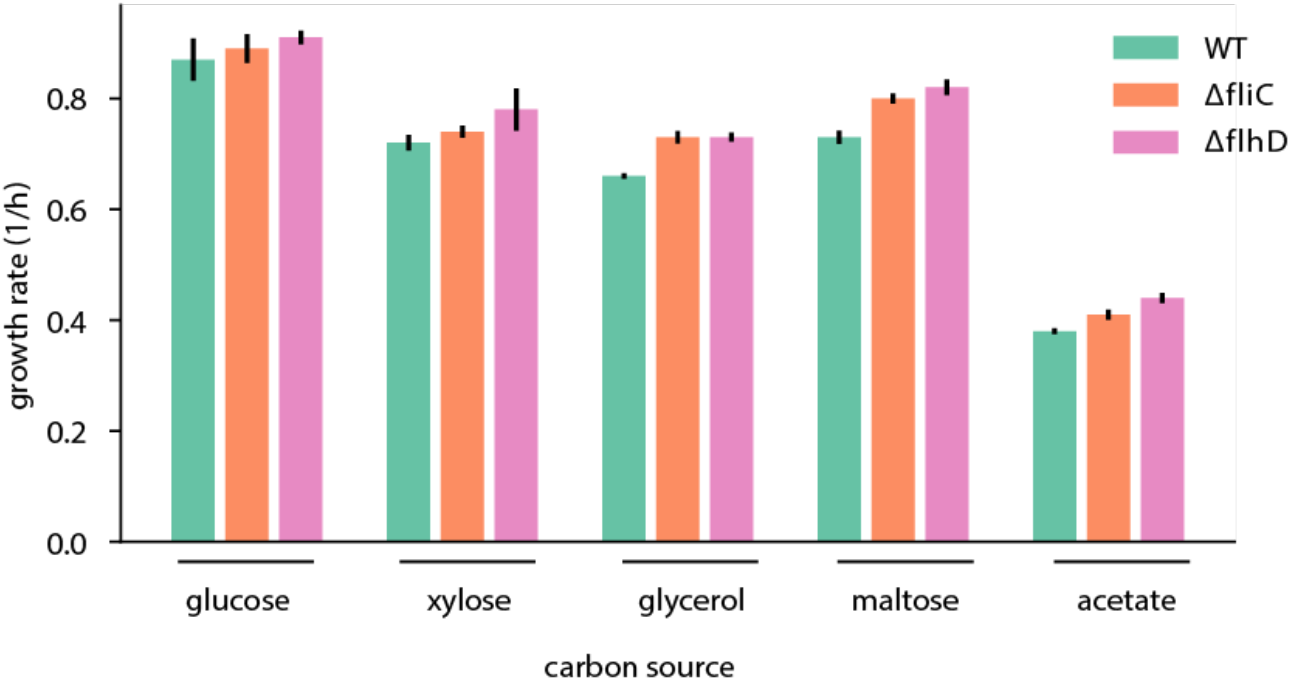
Balanced growth of deletion strains. Steady growth of WT and motility deletion strains *ΔfliC* and *ΔflhD* (trains listed in **Table S2).** Growth with 20mM glucose, 20mM xylose, 20mM glycerol, 20mM maltose, and 30mM acetate. Error bars show standard error based on N=3 biological repeats.

