## Supplementary Text for "Suboptimal proteome allocation during changing environments constrains bacterial response and growth recovery"

### Supplementary Information

Rohan Balakrishnan[1], Terence Hwa[1,2], Jonas Cremer[3]

#### TABLE OF CONTENTS

Supplementary Text .......... 2

1. Experimental Method 2
1.1 Strain information 2
1.1.1 Construction of deletion strains 2
1.2 Media and growth conditions 2
1.2.2 Lag-time quantification 3
1.2.3 Overexpression experiments 3
1.3 Sample analysis 3
1.3.1 qPCR measurements 3
1.3.2 RNA-Seq sampling and analysis 4
2 Modeling growth-kinetics during the shift 4
2.1 Growth on one nutrient source 5
2.2 Modeling nutrient consumption 7
2.3 Model parameters for growth on one carbon source 8
2.4 Modeling growth-transitions 8
2.5 Model parameters 10
2.6 The delaying effect of non-needed protein expression on growth-transition 11
2.7 Variation of pre-shift expression of required proteins. 11
2.8 Ribosome content 12

Supplementary Tables .......... 12

Table S1: Primers used in this study 12
Table S2: Strains used in this study 13
Table S3: Model parameters 13

Reference .......... 14

### SUPPLEMENTARY TEXT

#### 1. Experimental Method

##### 1.1 Strain information

The wild-type strain we use is the extensively characterized *E. coli* K-12 strain NCM3722(1, 2). This strain was also used as the parent for the construction oa all the strains used in this study.

###### 1.1.1 Construction of deletion strains

To obtain the *fliC* and flhD deletion strains, the corresponding KO strains JW1908 from the Keio collection(3) was used and the deletion was subsequently moved into the NCM3722 strain by phage P1*vir* transduction, yielding strains GE029 and NQ1225. All strains are listed in Table S2.

##### 1.2 Media and growth conditions

All growth media used in this study were based on the MOPS-buffered minimal medium used by Cayley et al. with slight modifications(4). 20 mM $NH_4Cl$ was provided as nitrogen source. Trace micronutrients were added as indicated in (4). For growth measurements in minimum medium, one of the followings was used as the primary carbon source: 20 mM glucose, 30 mM acetate, 20 mM glycerol, 20 mM sorbitol, 20mM succinate, 6 mM mannose, 4 mM mannose, 20 mM xylose. For shift experiments, cells were grown with 0.61mM glucose and the concentration of the second carbon source as stated before (e.g. 30mM acetate).

For the titratable strains (NQ1389 for *lacZ* and NQ1350 for *aceBA*), different concentrations of chlortetracycline (cTc) to induce expression and 15 ug/ml chloramphenicol and 50 ug/ml ampicillin to maintain the plasmid construct were additionally provided.
Cells were grown in a 37°C water bath shaker shaking at 250 rpm. To ensure balance growth, cells grew exponentially for at least 7 generations before starting measurements. We measured optical density at 600nm (OD600) using a UV-Vis spec. To obtain the growth rate of steadily growing cultures, OD600 data points within the range 0.04 to 0.4 (linear range of spectrophotometer) were obtained and fitted to an exponential growth curve. In addition, growth-curves and transitions were also quantified in a microplate-reader (200ul per well). A Tecan Spark microplate reader was used and absorbance (420nm) was measured every 7 min; incubation temperature was set to 37°C. Between measurements, plates were shaking at 132 rpm with an orbital amplitude. Wells loaded with only media (no culture) were used to reset absorbance values, and obtained absorbance values were subsequently adjusted to obtain OD600 values matching the values obtained with the UV-Vis spectrophotometer and a pathlength of 1cm. Obtained growth rates for glass-tube cultures and incubation in the microplate reader are highly comparable ($< \pm 5\%$). Growth-rate measurements were repeated 2-3 times as indicated in the figure captions.

##### 1.2.2 Lag-time quantification

To quantify lag-times, we first fit exponential growth behavior to the two steady growth phases (growth on glucose and growth on acetate) using OD600 ranges (0.04…0.15) and (0.3…0.4) for growth on glucose and acetate, respectively. Plateau levels (no change in OD600) were then determined by hand (OD value which first derivatives from the exponential growth on glucose), and the times $t_{pl,glucose}$ and $t_{pl,acetate}$ were the exponential curves match the plateau levels were calculated. The lag-time is the difference of these times, $t_{lag} = t_{pl,acetate} - t_{pl,glucose}$. Times were readjusted before plotting ($t \rightarrow t - t_{pl,glucose}$, such that time=0 corresponds to the time were the exponential growth on glucose hits the plateau level (beginning of shift). Lag-time estimation is further described in **Figure S1A**. Experiments to quantify lag-times were repeated at least 3 times as indicated in the figure captions.

##### 1.2.3 Overexpression experiments

Overnight pre-cultures (OD600~0.5) were diluted to a starting OD600 of ~0.02. To ensure a simultaneous entry into the shift phase (time and density glucose runs out) for different inducer levels, a main culture was prepared in an Erlenmeyer flask and after 1 hour of incubation this main culture was split into different cultures (glass-tubes with 6ml culture each). A different amount of inducer stock (cTc) was then added for the different cultures at the moment of glucose runout. To ensure the addition of the inducer exactly at this time, a control culturing starting with slightly higher cell density was run in addition which entered the shift approximately 30min earlier. The observed density value at the shift was used as the indicator when to add the inducer levels to the main cultures. Obtained lag-times were highly reproducible for inducer levels up to $5ng/ml$. For higher inducer levels (lag-times >5h), variation was higher, presumably because the slightly late addition of inducer levels leads to small variation in bottleneck enzymes which can have strong consequences over long times.

#### 1.3 Sample analysis

##### 1.3.1 qPCR measurements

RNA was extracted using the TRIzol (Thermo Fisher) method combined with a column-based purification step. In detail, 2ml culture was collected 30min after the shift (30min after glucose run out) and spined down. Pellet was immediately resuspended in 250ul TMN buffer (10 mM Tris pH 8, 10 mM $MgCl_2$ and 60 mM $NH_4Cl$) and thoroughly mixed with 250ul Trizol. After 5 min incubation, 50ul chloroform was added and, after mixing and another minute incubation, the sample was centrifuged for 10min at 4C. The clear phase was collected and the obtained RNA was immediately washed using a RNA purification kit (Zymo Research RNA Clean & Concentrator-5, following the instructions). To remove plasmid DNA, a DNAase digestion step on the purification colums was added following the instructions. To quantify transcription, a two-step qPCR approach was chosen. Reverse transcription was first run with random hexaprimers to obtain cDNA (azura genomics, AzuraFlex cDNA Synthesis Kit following instructions with 1ul of RNA sample). Real time PCR reactions (10ul final volume) were prepared using a Sybr Green master mix (Biorad SsoAdvanced Universal SYBR@ Green Supermix), following the instructions and using 10x diluted cDNA sample. Primer sets for 16S RNA and the genes *aceB*, *aceA*, and *lacZ* were used as listed in **Table S1**. 300nmol primer concentration were used, standard curves confirmed the linearity for each of the four primer sets chosen, and melting curves confirmed selectivity of the reaction. The PCR was run in a Biorad CFX 384 instrument, with the protocol following the reagent instructions provided with the master mix. To calculate relative expression levels, obtained gene expression levels were normalized to the 16S RNA levels measured from the same cDNA sample. The expression level of, for example, *aceB* was calculated as $2^{-(c_{w,aceB}-c_{W,16S})}$. To compare the changes in expression for different inducer levels, values were in addition normalized by the expression level obtained for 0 inducer level. Measurements were repeated at least 3 times starting with different cultures (biological repeats).

##### 1.3.2 RNA-Seq sampling and analysis

Steady state cultures were grown to $OD_{600}$ of 0.5. 10 ml samples were spun and cells were resuspended in 200ul TMN buffer (10 mM Tris pH 8, 10 mM $MgCl_2$ and 60 mM $NH_4Cl$). RNA was extracted using 200ul TRIzol reagent followed by ethanol precipitation. Ribosomal RNA was removed using the Ribo-Zero kit (Illumina) and barcoded RNA-seq libraries were generated using the TruSeq stranded kits (Illumina) as per the vendor's protocol. The libraries were sequenced using Illumina's Hiseq4000 platform. Typically, around 20 million reads were obtained per sample, except for the WT sample 5 minutes post shift, which had 5 million reads. Reads were de-multiplexed and aligned to the *E.coli* MG1655 U00096.3 genome using bowtie v2.2.6(5). Read counts were obtained using Python HTSeq-count (HTSeq v0.6.1p2)(6).

#### 2 Modeling growth-kinetics during the shift

Modeling growth-transitions is challenging because one needs to analyze how core growth-processes change with current growth-conditions which depends on the metabolic state of the cell. More recently, several modeling approaches have been formulated to overcome these challenges and to investigate growth transitions in different changing environments, including defined down- and up-shifts to carbon sources supplying faster or slower growth. These models consider the growth-conditions during growth-transitions focusing on the link to observed steady state growth behavior, parameterizing for example the fluxes such that they merge with steady state conditions(7, 8). These approaches extended the logic of growth-laws from steady state considerations to also describe the growth kinetics during the transition. To explicitly analyze the promoting effect of required metabolic proteins and the inhibiting effect of non-required genes on growth-transitions, we built on these models but chose a more explicit approach and specifically considered the expression of required and non-required enzymes during the shift. Model logic and details are provided in the following. Model results are shown in **Figs. 3 & S4**.

#### 2.1 Growth on one nutrient source

To a model for growth-kinetics we first introduce a simpler model for the growth on one carbon source. The model focuses on novel protein synthesis, the most resource demanding cellular component, and considers the allocation of ribosome activity to different protein classes as introduced in **Fig. S3.** We specifically build on the rationale introduced in Scott et al(9) to first consider balanced growth on one carbon source. Proteins are synthesized by ribosomes and the increase of total protein mass in the culture (variable $M$) depends on the number of ribosomes $N_{Rb}$ in the culture and their average translation speed $k_{Rb}$ (how many new amino-acids a ribosome is synthesizing per time):

$$\frac{dM}{dt} = k_{Rb} N_{Rb}$$

For convenience, we measure protein mass in numbers of amino acids such that no extra conversion factor is needed. Instead of the number of ribosomes we can also consider their mass (variable $M_R$) writing:

$$\frac{dM}{dt} = \gamma M_R$$

With the translation efficiency $\gamma_{Rb} \equiv k_{Rb}/m_{Rb}$ being a rate (unit 1/time) and $m_R = 7459AA$ the number of amino acids per ribosome.

The increase of ribosomal mass depends in turn on how fast proteins are synthesized, and to which extend the cell is allocation translation resources towards the synthesis of novel ribosomes (the thickness of arrow 3 in **Fig. S3A**). We write:

$$\frac{dM_R}{dt} = \alpha_{Rb} \frac{dM}{dt}$$

and call $\alpha_{Rb}$, a number between 0 and 1, the allocation coefficient towards ribosome synthesis.

To understand how biomass is increasing, we next have to consider translation in more detail. Notably, the translation efficiency $\gamma$ is not a constant rate but it is changing with the tRNA- precursor concentrations ribosomes encounter within the cells; ribosomes rely on a sufficiently high concentration of charged tRNA to work efficiently. Let us thus introduce a variable $p$ describing precursor concentrations within the cell. If $p$ falls to low levels, translation-speeds and thus biomass accumulation drops. In a first proxy, this can be described by a simple Michaelis-Menten relation:

$$\gamma = \gamma(p) = \gamma_{max} \frac{p}{p + p_0}$$

The Michaelis-Menten constant $p_0$ can be taken from measurements characterizing how translation falls with tRNA concentrations. To calculate precursor concentrations $p$ we consider the total precursor mass $M_{pc}$ and compare it to the total protein mass $M$ in the culture in the cell as total precursor mass $M_{pc}$ in the culture per total protein mass in the culture: $p \equiv M_{pc}/M$ (a conversion can give precursors per cell volume or dry mass). To investigate how the precursor concentrations change (over time and depending on parameters), we consider the change of total mass of precursors:

$$\frac{dM_{pc}}{dt} = J_{pc,in} - \frac{dM}{dt}$$

Precursor mass is given by a balance of novel precursor synthesis (flux $J_{in}$) and the utilization by the ribosomes increasing the total protein mass ($dM/dt$). The supply of precursors depends on the joint activity of metabolic enzymes which take up nutrients and make sure they are converted to amino acids, energy, and finally charged tRNAs (the precursors ribosomes need to grow). This is a complex process which we describe here in first order by jointly considering all metabolic enzymes as one major protein class and a simple 1st order reaction, writing $J_{pc,in} = k_{Mb} M_{Mb}$. $M_{Mb} is$ the mass of the metabolic protein class. $k_{Mb}$ is an effective rate describing how fast these proteins generate pre-cursors which we here call *metabolic efficiency*. For the precursor mass in the culture, we thus have:

$$\frac{dM_{pc}}{dt} = k_{Mb} \ M_{Mb} - \frac{dM}{dt}$$

Similarly as for the ribosomes, the increase of metabolic proteins depends on the allocation of translation activity towards these enzymes. We write:

$$\frac{dM_{Mb}}{dt} = \alpha_{Mb} \frac{dM}{dt}$$

Here, $\alpha_{Mb}$ denotes the allocation parameter towards the pool of metabolic enzymes. Notably, ribosomes can only translate one protein at a time, which leads to the overall constraint that the allocation parameters need to add up to 1. In the simplest case, assuming the cell only needs to generate metabolic enzymes and ribosomes:

$$\alpha_{Mb} + \alpha_{Rb} = 1.$$

Since cells also need to synthesize many other enzymes needed for growth which are not directly involved in precursor supply and translation(10), we extend this to:

$$\alpha_{Mb} + \alpha_{Rb} + \alpha_0 = 1$$

with $\alpha_0$ denoting the allocation towards synthesizing all other proteins. All three protein classes and the allocation of protein synthesis towards those are illustrated in **Fig. S3A**. The constraint is described by the relative thickness of the 3 arrows.

With this formulation we can analyze the balanced exponential growth which emerges:

$$\frac{1}{M}\frac{dM}{dt} = \lambda = constant$$

when the cellular concentrations and fractions are not changing over time, $\frac{dp}{dt} = \frac{1}{M}\frac{dM_{Rb}}{dt} = \frac{1}{M}\frac{dM_{Mb}}{dt} = 0$ and $\frac{M_{Rb}}{M} = \alpha_{Rb}$ and $\frac{M_{Mb}}{M} = \alpha_{Mb}$. The growth rate $\lambda$ depends on the rates (translation efficiency $\gamma_{max}$ and metabolic efficiency $k_{Mb}$), as well as the allocation parameters $\alpha_{Rb}$ and $\alpha_{Mb}$ . It can be calculated by the solution of a quadratic equation. Notably, the cell appears to control novel ribosome synthesis and adjusts the allocation parameter $\alpha_{Rb}$ towards optimizing growth-rates: ribosome synthesis is regulated such that precursor levels are optimal, and ribosomes can translate close to full speed(2). For our purpose this means that we know the allocation towards ribosomes for steady growth on glucose (or other carbon sources) and we can parametrize the model for steady state growth and subsequently extend this description to analyze growth-transitions.

#### 2.2 Modeling nutrient consumption

Up to now we have not explicitly considered nutrient consumption but assumed that precursor influx via the metabolic enzymes is described by a constant metabolic efficiency $k_{Mb}$. To consider nutrient availability, a crucial step towards describing shifts, we model the metabolic efficiency $k_{Mb}$ to be dependent on the nutrient concentration (say glucose, $n_{glu}$) in the culture:

$$k_{Mb}(n_{glu}) = k_{Mb,max}\frac{n_{glu}}{n_{glu} + K_{M,glu}}$$

$k_{Mb,max}$ denotes the maximum efficiency when nutrients are not limiting. $K_{m,glu}$ is the Monod constant for growth on glucose. To describe the change of nutrient availability, we consider the consumption of all nutrient molecules in the culture (nutrient mass $N_{glu}$ ) and write:

$$\frac{dN}{dt} = -k_{Mb}M_{Mb}/Y_{glu}$$

Here, the yield $Y_{glu}$ describes the conversion from nutrients (glucose) to precursors (charged tRNA). To obtain the yield in units of amino-acids one can take measured yield value in units of dry weight per nutrient weight and then convert assuming a fraction of 60% dry-weight content being proteins made up of amino acids with an average weight of $118.9 g/mol$.

#### 2.3 Model parameters for growth on one carbon source

To further parametrize the model, we can take translation speeds from in-vivo measurements, and the allocation parameters from steady state growth analysis across growth-conditions (see above). With that, the metabolic efficiency $k_{Mb}$ is the only parameter remaining and we adjust this parameter such that the emerging growth-rate matches the one experimentally observed for growth on glucose. Parameters are provided in **Table S3**. The resulting dynamics is shown in **Fig. S4A-E** (brown dashed lines which initially follow the black lines) for a culture starting with a glucose concentration of 0.6mM. Cells grow steadily with a constant growth-rate before glucose runs out, precursor concentrations drop, and cell-growth stops.

#### 2.4 Modeling growth-transitions

To describe growth-transition from growth on glucose towards growth on acetate we build on the formulation for growth on glucose described before and consider how cells start to express the required enzymes (e.g. *AceB*) to replenish precursors once glucose is depleted. To do this, we introduce a second class of metabolic enzymes with mass $M_{Mb,ace}$ (**Fig. S3B**). Notably, this class of enzymes includes only those enzymes which are needed to ensure a recovery of the precursors influx (glyoxylate shunt and gluconeogenesis genes, **Fig. S1B**) and not those metabolic enzymes which are also needed for growth on glucose and thus already available and working (like TCA enzymes or enzymes of the respiratory chain to provide energy). $M_{Mb,ace}$describes thus a much smaller pool of proteins than what is eventually needed for balanced growth on acetate. Since we are interested in explaining lag-times until precursor levels recover, we here consider the requirements for those latter enzymes only indirectly by limiting the relative fraction of the enzyme class proteins ($M_{Mb,ace}/M$) to a maximum value (more below).

With the two metabolic fluxes the precursor turnover before and during the shifts is given by:

$$\frac{dM_{pc}}{dt} = k(n_{glu}) \cdot M_{Mb,glu} + k_{Mb,ace}(n_{ace}) \cdot M_{Mb,ace} - \frac{dM}{dt}$$

Nutrient concentrations in the culture changes accordingly:

$$\frac{dN_{glu}}{dt} = -k_{Mb,glu}(n_{glu})\, M_{Mb,glu}/Y_{glu}$$

$$\frac{dN_{ace}}{dt} = -k_{Mb,ace}(n_{ace})\, M_{Mb,ace}/Y_{ace}$$

Here, the metabolic efficiencies depend on the abundance of glucose and acetate respectively. As for growth on glucose alone, we model a Monod type relation with Monod constants $K_{M,glu}$ and $K_{M,ace}$ to describe how influx stops at low nutrient concentrations.

The synthesis of new metabolic enzymes depends on the allocation coefficients:

$$\frac{dM_{Mb,glu}}{dt} = \alpha_{Mb,glu}\frac{dM}{dt}$$

$$\frac{dM_{Mb,ace}}{dt} = \alpha_{Mb,ace}\frac{dM}{dt}$$

Depending on the availability of nutrients the cell is adjusting the allocation to these enzyme classes and to model growth-transitions we thus have to formulate relations describing how the allocation coefficients depend on the availability of both nutrient sources.

The allocation to enzymes required for growth on glucose is high during steady state growth on glucose but we assume that their expression reduces to lower levels when glucose-concentrations drop, we thus model:

$$\alpha_{Mb,glu} = \alpha_{Mb,glu,min} + \alpha_{Mb,glu,max}\left(\frac{n_{glu}}{n_{glu} + K_M}\right)$$

Here, $\alpha_{Mb,glu,max} + \alpha_{Mb,glu,min}$ is the same allocation coefficient we use to describe steady growth on glucose alone. In contrast, the allocation towards the required enzymes to recover precursor supply from acetate is only high once glucose runs out and this enzyme class is hardly expressed when glucose is still available. To account for this behavior, we use the following dependence on glucose concentrations:

$$\alpha_{Mb,ace} = \alpha_{Mb,ace,max}\left(1 - \frac{n_{glu}}{n_{glu} + K_M}\right)\left(1 - \frac{M_{ace}/M}{\frac{M_{ace}}{M} + \alpha_{Mb,ace,steady}}\right) + \alpha_{Mb,ace,preshift}\left(\frac{n_{glu}}{n_{glu} + K_M}\right)$$

That is, the synthesis of the novel metabolic enzymes required to provide precursors via acetate consumption is occurring by a certain fraction of translating ribosomes ($\alpha_{Mb,ace,max}$) once glucose is consumed. But this high rate falls again to the final steady state levels for growth on acetate once that fraction is reached. This limitation of the metabolic enzymes to a lower fraction allows us to indirectly consider that cells have to start synthesizing a broad class of metabolic proteins (like TCA cycle proteins) to grow once pre-cursor supply have been reestablished (and not only the glycolytic shunt and gluconeogenesis genes required immediately to rescue precursor influx). Finally, to study the role of pre-shift expression we also included an expression term $\alpha_{Mb,ace,pre-shift}$which describes expression when glucose is still abundant (in the reference condition this constant is 0).

As previously, the growth-kinetics is described by how the ribosomes synthesize new biomass:

$$\frac{dM}{dt} = \gamma(p)\ M_R$$

Translation and thus growth depends on precursor levels which in turn depend on the abundance and activity of the metabolic activities.

#### 2.5 Model parameters

We modeled growth transitions for a reference parameter set listed in **Table S3**. Here we provide further context.

To describe nutrient uptake towards precursor synthesis we used yield values ($Y_{glu}, Y_{ace}$) and Monod constants ($K_{m,glu}, K_{m,ace}$) known for growth on glucose and acetate. To determine the allocation parameters towards synthesis of the metabolic enzymes required to provide pre-cursers when growing on acetate ($\alpha_{Mb,ace,preshift}, \alpha_{Mb,ace,max}, \alpha_{Mb,ace,steady}$) we used the transcriptomics measurements we collected during the shift (Fig. S5). Given the fast turnover of mRNA this data provides a direct readout of the allocation behavior at different timepoints during the shift. We specifically calculated the relative mRNA fraction of all glyoxylate and gluconeogenesis genes which are required for the continuous influx of precursors when glucose runs out (**Fig. S1B**), and thus used 3% and 1% as reference values for $\alpha_{Mb,ace,max}$ and $\alpha_{Mb,ace,steady}$ respectively. We initially neglected pre-shift expression levels as those are very low, $\alpha_{Mb,ace,preshift} = 0$.

With the allocation parameters defined, only one fitting parameter remains, the rate $k_{Mb,ace}$describing how fast metabolic enzymes recover pre-cursors from acetate. We adjusted this rate such that the lag-time of the modeled growth-transition approximately matches the lag-time in the experiments (3.5h for the shift of WT cells from growth in glucose to growth acetate, **Fig. 1A**). With these parameters, the post-shift growth which emerges also resembles the growth-rate observed during the experiments. The simulated growth-transition for this reference parameter set is shown in **Fig. S4A-E** (black lines). Growth is fast in glucose, then stops temporarily and after a lag growth recovers by using acetate.

#### 2.6 The delaying effect of non-needed protein expression on growth-transition

With the formulated model and the given parameters, we can now investigate how growth-transitions are changing when the cell is allocating varying fractions of its translation activity during the shift to the required metabolic enzymes. Mathematically, this means varying the allocation parameter $\alpha_{Mb,ace,max}$. The results are shown in **Fig. S5F-J**. The lag-time falls strongly with a higher allocation towards the required enzymes, with the reciprocal relation shown in **Fig. 3B**. As discussed in **Fig. S5**, different allocations lead to varying drops of precursor levels during the shift which changes the ability to recover growth.

#### 2.7 Variation of pre-shift expression of required proteins.

Our transcription results show that the enzymes required to recover growth once glucose runs out are hardly expressed when glucose is still abundant (Fig. S6), suggesting that pre-shift expression levels do not play a major role in shaping lag-times for WT *E. coli* cells for these growth-conditions. But a higher pre-shift expression might occur in specific growth conditions and has been shown to support faster transition in a synthetic strain were pre-expression of the required genes *aceB/aceA* was controlled by an inducer construct(10). To investigate the role of pre-shift expression compared to expression during the shift we determined the emerging lag-times when both, the allocation before the shift ($\alpha_{Mb,ace,preshift}$) and during the shift ($\alpha_{Mb,ace,max}$) are varied (Fig. N1).

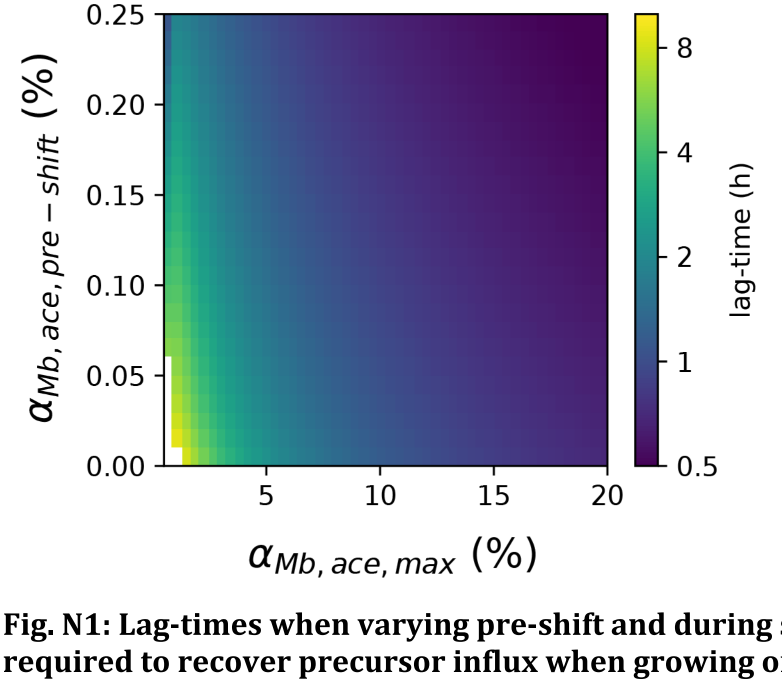


**Fig. N1: Lag-times when varying pre-shift and during shift expression of the genes required to recover precursor influx when growing on acetate.** We varied both allocation parameters $\alpha_{Mb,ace,preshift}$ and $\alpha_{Mb,ace,max}$ independently. Pre-shift expression of the enzymes can clearly lead to a faster lag-times. Notably, however, lag-times can also be short without pre-shift expression if cells response to the shift with a strong expression of required genes instead of the expression of other genes. The previously described tradeoff between fast pre-shift growth and fast shift thus appears to be a secondary tradeoff effect important when there are long lag-times in the first place because cells response by the expression of non-required enzymes.

##### 2.8 Ribosome content

Up to now, we have not explicitly modeled the change of ribosomes synthesis during the shift. An explicit consideration of ribosome synthesis is not required to investigate the origin of lag-times since ribosomes are highly abundant at the moment glucose runs out and they do not contribute to a bottleneck in precursor influx. And since ribosome synthesis abruptly stops at the shift (**Fig. S5G**) we do not have to consider the role of novel ribosome synthesis as a process competing for the same limited precursors required to synthesize the required enzymes to recover metabolic flux. Indeed, when explicitly modeling the change of novel ribosome synthesis lag-times do not change (data not shown). Particularly we modeled that ribosomes are only synthesized when precursor levels are above a threshold value, in line with what is currently known about the control of ribosome synthesis mediated by central regulators like the alarmone ppGpp. The allocation we modeled is:

$$\alpha_{Rb}(p) = \alpha_{Rb,steady} \cdot p/(p + p_0).$$

### SUPPLEMENTARY TABLES

##### Table S1: Primers used in this study

| Name | Sequence | Notes |
|---|---|---|
| aceA-qpcr-F | GCGTTGGGAAGGCATTACTCGC | qPCR |
| aceA-qpcr-R | GCCTGACCGCCAGTCAGTGC | qPCR |
| aceB-qpcr-F | GGCAACAACAACCGATGAACTGGC | qPCR |
| aceB-qpcr-R | GCTCAGTCAGAAATTCTACCGC | qPCR |
| lacZ-qpcr-F | GGCAATTTAACCGCCAGTCAG | qPCR |
| lacZ-qpcr-R | GTGCACGGGTGAACTGATC | qPCR |

| | | |
|---|---|---|
| 16S-qpcr-F | CCTAGGCGACGATCCCTAGC | qPCR |
| 16S-qpcr-R | CATACACGCGGCATGGCTGC | qPCR |
| fliC-F | CAGGGTTGACGGCGATTGAG | Confirmation KO |
| fliC-R | CAATTTGGCGTTGCCGTCAG | Confirmation KO |
| k1 | CAGTCATAGCCGAATAGCCT | Confirmation KO(1) |
| k2 | CGGTGCCCTGAATGAACTGC | Confirmation KO (1) |
| kt | CGGCCACAGTCGATGAATCC | Confirmation KO (1) |

**Table S2: Strains used in this study**

| Strain name | Description | Details (genotype, plasmid) | Ref |
|---|---|---|---|
| NCM3722 | Wild type (K12), genetic background for all strains used in this study | | (1) |
| NQ1389 | *lacZ* over-expression strain | Ptet-tetR on pZA31; Ptetstab-lacZ on pZE1 | (11) |
| NQ1350 | *aceBA* induction | aceBA promotor controlled by Ptet | (10) |
| GE029 | *ΔfliC* | | This study |
| NQ1225 | *ΔflhD* | | (12) |

**Table S3: Model parameters**

| Symbol | Description | Value |
|---|---|---|
| $\alpha_{Rb,glu}$ | Allocation to translation (ribosomes) | 0.2 |
| $\alpha_{Mb,glu}$ | Allocation to metabolic proteins | 0.45 |
| $k_{Mb,glu}$ | Metabolic efficiency glucose | $2.4\ 1/h$ |

| | | |
|---|---|---|
| $Y_{glu}$ | Yield in glucose | $0.377\ OD_{600}/mmol$ |
| $K_{M,glu}$ | Monod constant, glucose | $5\mu mol$ |
| $\alpha_{Rb,ace,steady}$ | Allocation to translation (ribosomes) | 0.2 |
| $\alpha_{Mb,ace,max}$ | Allocation to required enzymes at shift (or varied) | 0.03 |
| $\alpha_{Mb,ace,steady}$ | Allocation to required enzymes at shift at steady growth | 0.01 |
| $k_{Mb,ace}$ | precursor-rate for bottleneck enzymes (when other metabolic enzymes are ready) | $30\ 1/h$ |
| $Y_{ace}$ | Yield in acetate | $Y_{glu}/2$ |
| $K_{M,ace}$ | Monod constant | $5\mu mol$ |
| $Type\ equation$ | Conversion between OD and AA. Based on known conversion between OD and biomass, 0.63% of dry mass being proteins. | $7.69\ 10^{-18}\ OD_{600} \cdot ml/AA$ |
| $f_{Rb}$ | fraction active ribosomes | 0.65 |
| $k_{Rb}$ | Max. translation elongation rate | $20\ 1/s$ |
| $p_0$ | Michaelis Menten constant for translation. In units of charged tRNA per mass amino acids. | 0.026 |
